# Socioeconomic Resources are Associated with Distributed Alterations of the Brain’s Intrinsic Functional Architecture in Youth

**DOI:** 10.1101/2022.06.07.495160

**Authors:** Chandra Sripada, Arianna Gard, Mike Angstadt, Aman Taxali, Tristan Greathouse, Katherine McCurry, Luke W. Hyde, Alexander Weigard, Peter Walczyk, Mary Heitzeg

**Affiliations:** Department of Psychiatry, University of Michigan, Ann Arbor; Department of Psychology and Neuroscience and Cognitive Neuroscience Program, University of Maryland, College Park; Department of Psychology and Survey Research Center at the Institute for Social Research, University of Michigan, Ann Arbor

**Keywords:** socioeconomic resources, socioeconomic status, parental education, household income, neighborhood disadvantage, resting state fMRI, functional connectivity, connectomics, Adolescent Brain Cognitive Development (ABCD) Study, predictive modeling, intrinsic connectivity networks, neurodevelopment

## Abstract

Little is known about how exposure to limited socioeconomic resources (SER) in childhood gets “under the skin” to shape brain development, especially using rigorous whole-brain multivariate methods in large, adequately powered samples. The present study examined resting state functional connectivity patterns from 5,821 youth in the Adolescent Brain Cognitive Development (ABCD) study, employing multivariate methods across three levels: whole-brain, network-wise, and connection-wise. Across all three levels, SER was associated with widespread alterations across the connectome. However, critically, we found that parental education was the primary driver of neural associations of SER. These parental education associations were robust to additional tests for confounding by head motion, and they exhibited notable concentrations in somatosensory and subcortical regions. Moreover, parental education associations with the developing connectome were partially accounted for by home enrichment activities, child’s cognitive abilities, and child’s grades, indicating interwoven links between parental education, child stimulation, and child cognitive performance. These results add a new data-driven, multivariate perspective on links between household SER and the child’s developing functional connectome.

## INTRODUCTION

Childhood socioeconomic resources (SER), reflecting a combination of parental education and economic resources within home and neighborhood contexts, shape adult outcomes, particularly in economic (e.g., earnings, employment), educational (e.g., cognitive skills, college completion), and physical and mental health domains (1–3). Inequalities in access to SER is larger in the United States than other industrialized countries (4) and have been growing over time (5, 6), and thus SER-related inequalities are likely to have wide-ranging impacts on wellbeing across the life course (7). This implication has encouraged neuroscientists to investigate pathways through which SER influences the developing brain (8, 9). Yet our current understanding of these pathways remains highly incomplete, particularly during critical developmental windows such as early adolescence, marked by extensive neural reorganization (10) and when many serious psychosocial challenges (e.g., problems in interpersonal, academic, and mental health domains) first emerge (11).

The human brain is organized as a complex network (12, 13), with interconnections among regions implicated in diverse cognitive and socioemotional functions (14). Task-free “resting state” functional magnetic resonance imaging (fMRI) uses coherence in spontaneous activity across brain regions to yield maps of functional connectivity patterns (15), which, in turn, can be linked to individual difference variables such as such cognition, personality traits, or psychopathology (16).

Previous studies of the impact of SER on resting state functional connectivity patterns have mostly relied on region-specific approaches that focus on individual connections (e.g., amygdala-ventromedial prefrontal connectivity) (17), requiring strong a priori knowledge about which connections are (and are not) implicated. There is, however, convergent evidence that characteristics of social, psychological, and clinical interest often involve distributed and wide-ranging changes at tens of thousands of connections distributed across the entire brain (18), rather than focal changes involving individual pairs of regions. Additionally, previous studies often used small samples consisting of tens to hundreds of subjects. Recent widely discussed results (19) demonstrate that these studies are liable to produce spurious findings, and several thousand subjects are typically needed to derive statistically reliable conclusions. At the present time, however, no previous studies have investigated SER-associated functional connectivity patterns using multivariate methods across tens of thousands of brain connections in large, adequately powered samples.

Additionally, SER is a multi-dimensional construct that incorporates features of parental education, household income, and neighborhood disadvantage (20). Different components of SER likely implicate different underlying environmental mechanisms (e.g., maternal education may impact cognitive stimulation in the home, whereas neighborhood disadvantage may operate through school quality). Understanding which component(s) of SER sculpt the developing brain is critical for designing targeted interventions and for informing housing, school, and redistributive policies (21, 22). But the unique effects of each dimension of SER in shaping brain-wide connectivity patterns remains unclear.

To address these gaps in understanding, we leveraged the Adolescent Brain and Cognitive Development (ABCD) Study (23, 24), a population-based study of 11,875 9- and 10-year-olds from 22 sites across the United States with substantial sociodemographic diversity (25). ABCD is the largest developmental neuroimaging study ever undertaken, providing a unique opportunity to study how SER shapes connectivity patterns of the developing brain. To convergently establish results at multiple levels of analysis, we employed three complementary multivariate methods: a whole-brain approach (multivariate predictive modeling) (26); a network-wise approach (network contingency analysis) (27–29), and a connection-wise approach (quantile-quantile modeling) (30).

These analyses jointly indicated that a combination of all the three dimensions of SER (i.e., parental education, household income-to-needs, and neighborhood disadvantage) was associated with widespread individual variation in connectivity across the entire brain, with significant effects observed in 77 out of 120 “cells”, i.e., sets of connections linking pairs of large-scale brain networks. In additional analyses that dissected the unique contribution of individual components of SER, we found the most potent associations with parental education, even after controlling for the contributions of household income-to-needs and neighborhood disadvantage. Moreover, functional connectivity patterns associated with parental education were uniquely concentrated in sensorimotor and subcortical networks. Our results may help to illuminate why SER is associated with a variety of outcomes across the life course, while also highlighting the need for more research to explore proximal biopsychosocial mechanisms of SER-connectome associations.

## RESULTS

### 1. Across three levels of analysis (i.e., whole-brain, network-wise, and connection-wise), socioeconomic resources are associated with large, brain-wide changes in functional connectivity

#### Whole-Brain-Level Analysis

We built and assessed multivariate predictive models for socioeconomic resource (SER) scores using a leave-one-site-out cross-validation approach. At each fold of the cross-validation, we trained a multivariate predictive model to use individual differences in brain connectivity patterns to predict SER scores. We then applied the trained model to brain connectivity data from subjects at the held-out site, yielding predictions of their SER, and we repeated this sequence with each site held out once. We accounted for nuisance covariates (youth sex assigned at birth, age, self-reported race-ethnicity [a social construct linked to disparities in access to resources due to historical and present structural discrimination], and head motion) by applying regression coefficients for covariates learned in the train data to covariates in the test data, thus preserving complete independence between train and test datasets. We found that the correlation between actual versus predicted SER, controlling for covariates and averaging across the 19 folds of the cross-validation, was 0.28 (Figure 1, upper left panel). That is, after accounting for covariates, brain connectivity patterns accounted for 9.0% of the variance in SER in held-out samples of youth (cross-validated r^2^). Cross-site generalizability was remarkably consistent: Correlations between predicted and actual scores were statistically significant in all 19 out of 19 held-out sites (all 19 site-specific permutation *p*-values < 0.0001; observed correlations were higher than all 10,000 correlations in the permutation distribution).

**Figure 1:**
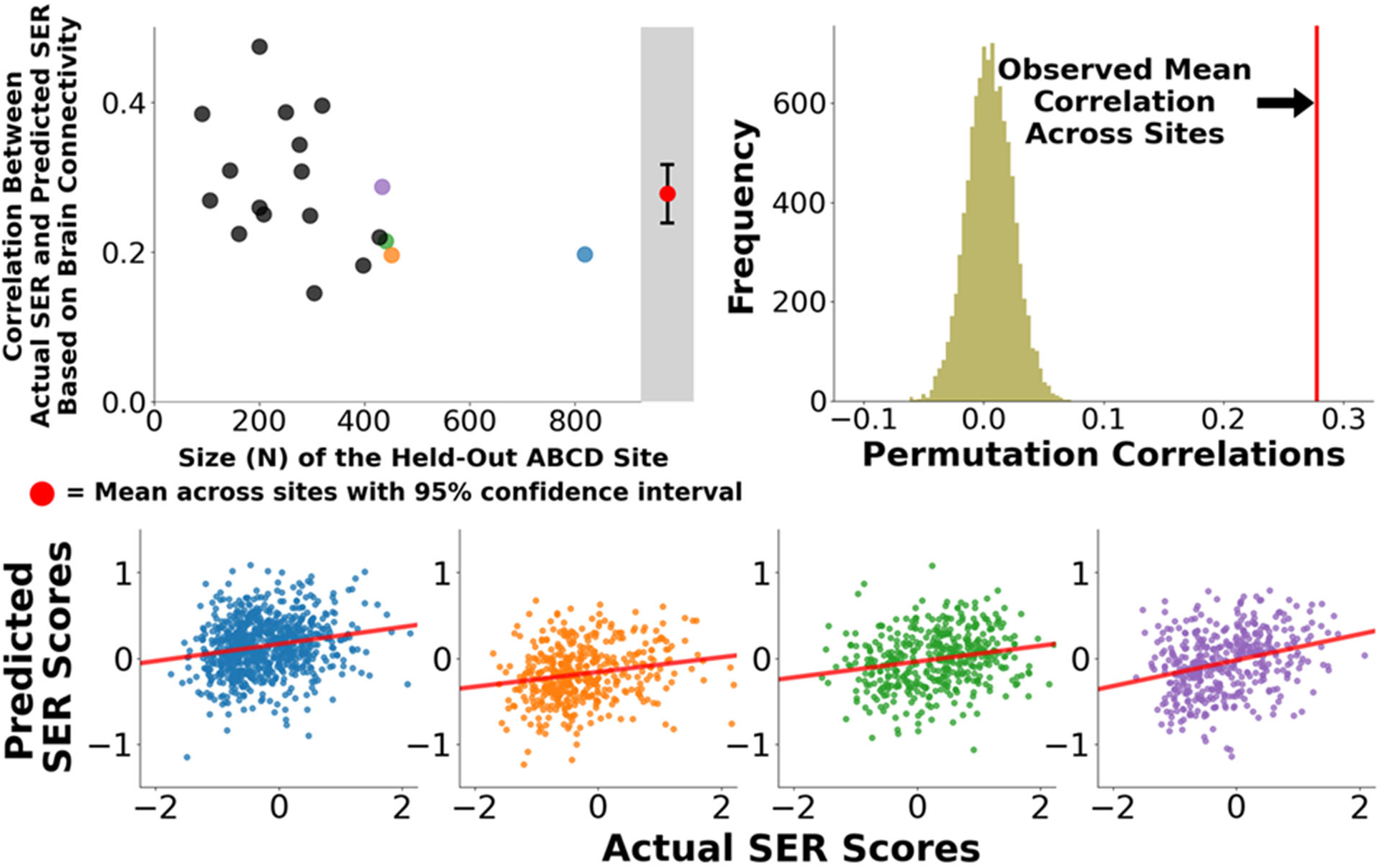
Correlations Between Actual Socioeconomic Resources Scores and Scores That Are Predicted Based on Whole-Brain Connectivity Patterns. We applied multivariate predictive models to 5,821 subjects at 19 sites to identify brain-wide connectivity patterns that are associated with SER scores. (Upper Left Panel) In leave-one-site-out cross-validation, functional connectivity patterns associated with SER scores generalized to 19 out of 19 held out sites. (Upper Right Panel) The overall mean correlation between observed SER scores and predicted SER scores (predicted exclusively from brain connectivity patterns) was 0.28, p_PERM_<0.0001 (observed correlation was higher than all 10,000 correlations in the permutation distribution). (Lower Panel) Scatter plots for four largest held-out sites (blue, orange, green, purple) show consistent performance at individual sites.

#### Network-Level Analysis

The human brain is organized into a number of large-scale networks (31) defining a set of “cells”, which are sets of connections linking pairs of large-scale networks. We performed network contingency analysis (NCA) (27–29), which identifies cells in which the count of connections associated with SER (controlling for covariates) exceeds the count expected by chance, established by non-parametric permutation tests. As shown in Figure 2, a total of 77 out of 120 cells exhibited significant associations with SER (FDR<0.05; shaded in the Figure 2), and these cells were notably widespread throughout the brain spanning all large-scale networks.

**Figure 2:**
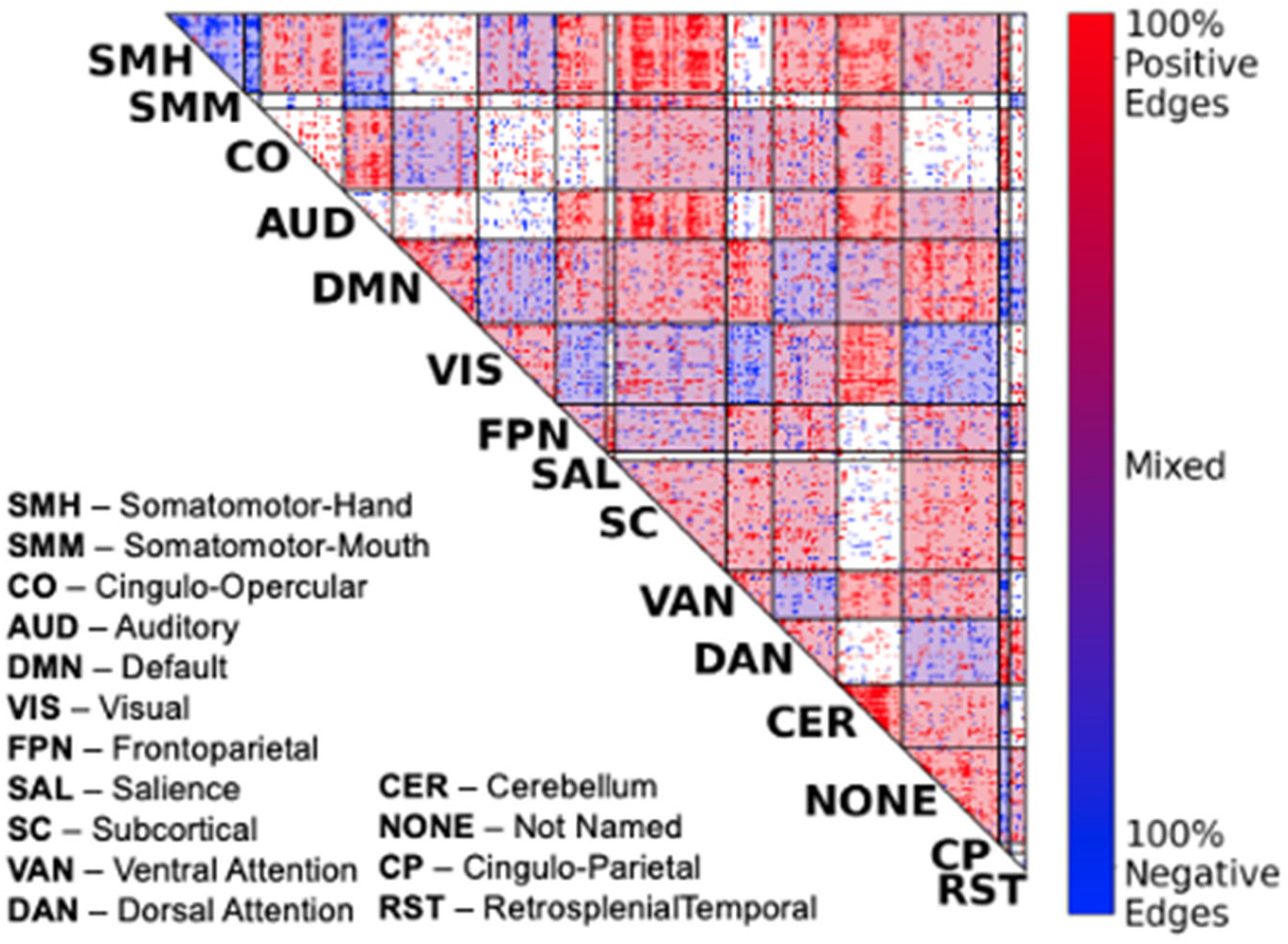
Network-to-Network Connections Exhibiting Significant Effects of Socioeconomic Resources. We performed network contingency analysis (NCA) which identifies cells (i.e., sets of connections linking pairs of large-scale networks) where the number of edges related to SER scores exceeds the number expected by chance. A total of 77 out of 120 cells exhibited significant effects of SER (FDR<0.05; shaded in the figure), and these cells were notably widespread throughout the brain.

#### Connection-Level Analysis

We additionally assessed associations with SER on a connection-by-connection basis using quantile-quantile modeling (30). We first calculated the p-value at each connection for the association between that connection and SER (controlling for covariates). We then rank ordered these p-values and plotted them against the rank-ordered distribution of p-values expected under the global null hypothesis, which was calculated with non-parametric permutation-based methods (Figure 3). If there is no association between SER and brain functional connections, this plot should follow the 45° line shown in in red in Figure 3. But the observed plot strongly deviated from this line. Moreover, “lift off”, where the observed distribution deviates from the 95% confidence interval of the null line, occurred very early and persisted through the range of the x-axis. This result is consistent with widespread associations of SER with most functional connections across the brain.

**Figure 3:**
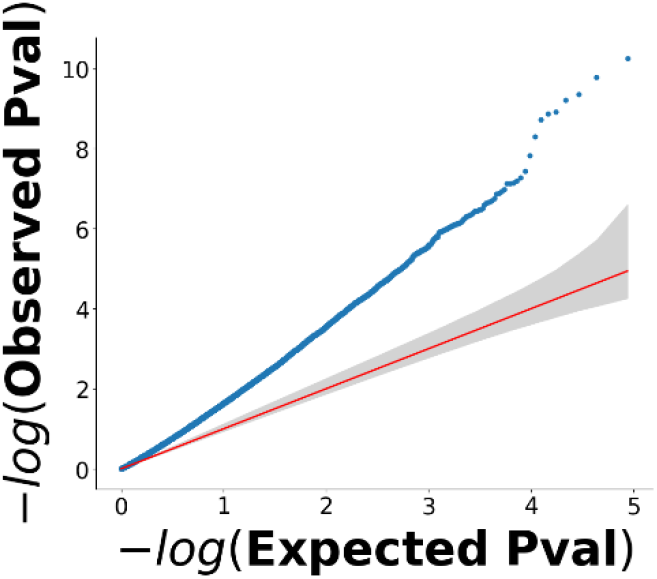
Quantile-Quantile Model of Effects of Socioeconomic Resources on Functional Connections of the Connectome. The red line and its associated 95% confidence interval (in gray) represent the global null hypothesis that household SER scores are unrelated to functional connections of children’s connectomes. The plot shows strong deviation of the observed data (blue line) from the red line with a pattern of early “lift-off”, in which the deviation occurs towards the left of the plot and is sustained throughout. This pattern is consistent with widespread, diffuse influences of SER scores throughout the connectome.

### 2. Parental education is the primary driver of brain-wide changes in functional connectivity, with household income-to-needs and neighborhood disadvantage having weaker (or absent) unique effects

The preceding results established strong and widespread influences of SER on children’s connectomes across whole-brain, network-wise, and connection-wise levels of analysis. We next sought to identify unique contributions of the three components of the SER variable: parental education, household income-to-needs, and neighborhood disadvantage. We thus repeated the preceding analyses, this time with one of these three dimensions of SER as the variable of interest and the other dimensions entered as additional covariates. We repeated these analyses three times in total with each SER dimension as the variable-of-interest.

Results showed that parental education (controlling for household income-to-needs and neighborhood disadvantage; left column in Figure 4) exhibited consistently strong effects at all three levels of analysis. Household income-to-needs (controlling for parental education and neighborhood disadvantage; middle column of Figure 4) showed modest but statistically significant effects in multivariate predictive modeling analysis, no significant cells in network analysis, and did not deviate from the null hypothesis line in quantile-quantile analysis. Neighborhood disadvantage (controlling for parental education and household income-to-needs; right column of Figure 4) did not show any statistically significant effects in any of the three analyses.

**Figure 4:**
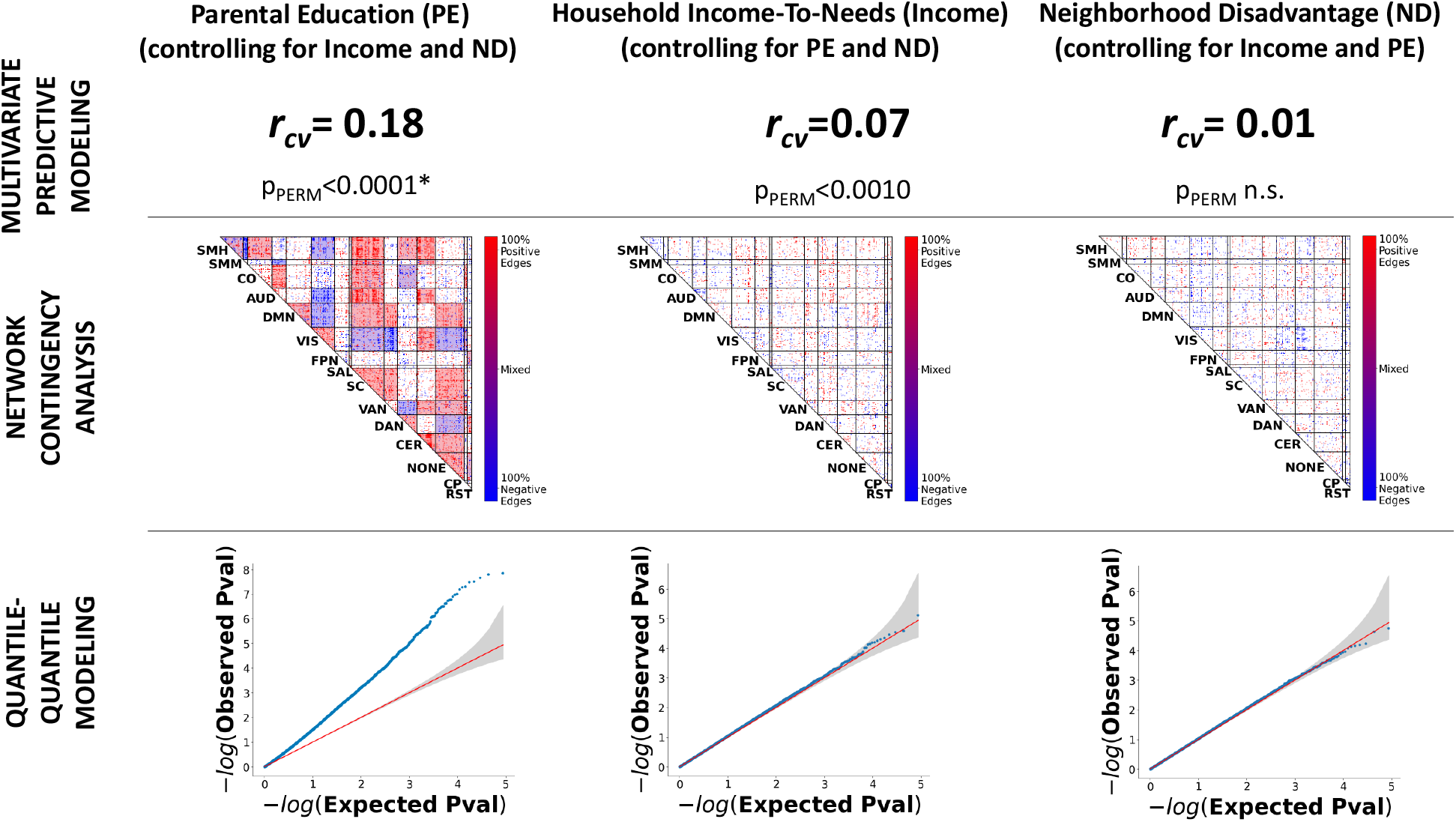
Parental education has widespread and unique effects on children’s resting state connectomes. We used whole-brain-level (top row), network-level (middle row), and connectionlevel (bottom row) methods to identify unique effects on children’s resting state connectomes of three SER variables: parental education (left column), household income-to-needs (middle column), and neighborhood disadvantage (right column), Across the three-levels of analysis, parental education (controlling for household income-to-needs and neighborhood disadvantage) demonstrated consistent strong effects. Household income-to-needs (controlling for parental education and neighborhood disadvantage) showed modest, statistically significant effects in multivariate predictive modeling analysis, but showed no significant cells in network analysis and did not deviate from the null hypothesis line in quantile-quantile analysis. Neighborhood disadvantage (controlling for parental education and household income-to-needs) did not show any statistically significant effects in all three analyses. r_cv_ = cross-validated out-of-sample correlation between actual scores and predicted scores using resting state connectivity data; * = observed correlation higher than all 10,000 correlations in the permutation distribution

### 3. Parental education connectomic associations were robust to additional tests for potential confounds including head motion

Given evidence that parental education is the primary driver of SER associations with the connectome, we subjected parental education to additional robustness tests in a low head motion subsample (mean FD<0.2mm, *N*=2844) to assess potential confounds associated with head motion, which has emerged a serious confound in youth functional connectivity studies. We focused on multivariate predictive modeling, which yields a single scalar performance measure (out-of-sample correlation), making it ideal for conducting these robustness tests. We found that in this low motion sample, out-of-sample correlations between observed and predicted parental education (both uncontrolled and controlling for household income-to-needs and neighborhood disadvantage) remained highly statistically significant (uncontrolled r_cv_=0.24; controlled r_cv_ = 0.12; p_PERMS_’s <0.0001; see Table S3).

Because correlations observed in testing samples are typically lower when models are trained on a smaller training subsample (32), we additionally performed resampling tests (see Supplement) to assess whether observed performance in the low motion subsample differs significantly from performance in randomly drawn subsamples of the same size. These resampling tests were non-significant (see Table S3), indicating that the observed out-of-sample parental education correlations in the actual low motion subsample does not statistically differ from other subsamples of the same size. Taken together, these results provide additional evidence that associations between parental education and children’s functional connectomes are unlikely to be due to confounding with head motion.

### 4. Parental education exhibits a sensorimotor/subcortical pattern that differentiates it from other youth phenotypes also associated with resting state connectivity

Given evidence of robust associations between parental education and resting state functional connectivity, we next compared the parental education connectivity pattern with corresponding connectivity patterns observed for general cognitive ability and the general factor of psychopathology, i.e., “P factor”, which were recently studied by our group in this same ABCD sample (28, 33). We focused on connectivity patterns observed with network contingency analysis, which is well-suited for localizing associations to sets of connections linking pairs of networks. Bubble graphs (Figure 5) capture the proportion of significant cells associated with each network for each variable. Consistent with our previous reports, the general factor of psychopathology implicated interconnections between the default mode network and a number of control networks (28), while general cognitive ability implicated highly distributed connectivity patterns involving all networks (33). Though there was some overlap across the three variables, parental education preferentially implicated somatosensory/subcortical networks, with especially prominent involvement of the visual and subcortical networks. Non-parametric tests of concentration showed that the proportion of significant cells within somatosensory/subcortical networks exceeded what is expected by chance (*p*_PERM_=0.003).

**Figure 5:**
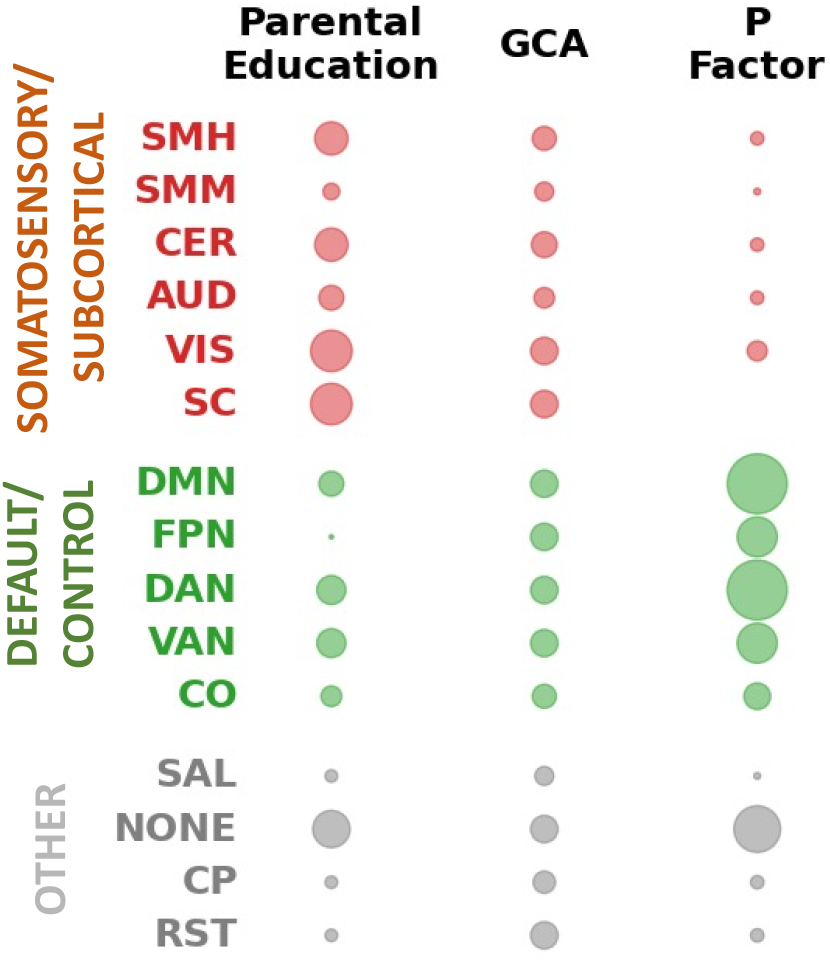
Parental Education Effects are Concentrated in Somatosensory/Subcortical Networks. We compared parental education connectivity patterns with corresponding patterns observed for general cognitive ability and the general factor of psychopathology (P Factor). The bubble graph captures the proportion of significant cells, i.e. sets of connections linking large-scale networks, associated with each network. The graph shows each variable has a distinct profile across the connectome, with parental education preferentially implicating somatosensory/subcortical networks.

### 5. Parental education’s associations with the functional connectome are related to home enrichment activities, child’s grades, and child’s cognitive abilities

Previous literature, reviewed in (34), suggests that parental education may impact children’s behavioral and cognitive outcomes through provision of greater cognitive stimulation and parenting behaviors that promote academic and social competence (e.g., warmth and support). To gain some initial insight into why parental education was associated with children’s brain connectivity patterns, we performed an exploratory analysis that accounted for potential additional explanatory variables: 1) parent-reported enrichment activities (e.g., having intellectual discussions, reading with the child); 2) child-reported family support (e.g., smiling at the child, discussing the child’s worries, providing support); 3) child-reported school support (e.g., availability of extracurricular activities, praise when the child does a good job); 4) child’s grades in school; and 5) child’s general cognitive ability. This analysis was performed in a subsample of 3,223 children for whom all the preceding variables were available. We focused on whole-brain predictive modeling, which affords a single dependent measure (out-of-sample brain-phenotype association) as well as quantitative bootstrap-based tests for attenuation of this association (see Supplement). Results showed that home enrichment activities, child’s grades, and child’s general cognitive ability each individually significantly attenuated the association between parental education and youth brain connectivity patterns (Table 1). In contrast, these associations were not meaningfully attenuated by controlling for family supportiveness and school supportiveness. In combination, 29% of the multivariate association between parental education and the functional connectome was accounted for by these five candidate explanatory variables.

**Table 1:** Associations Between Parental Education and Brain Connectivity Patterns Controlling for Aspects of the Family and School Environment and Child Characteristics. We quantified the attenuation of the relationship between parental education and brain connectivity patterns after controlling individually and jointly for six candidate explanatory variables. Overall, 1/3 of the multivariate association between parental education and brain connectivity patterns was accounted for by the six candidate explanatory variables. r_cv_ = cross-validated out-of-sample correlation between actual scores and predicted scores using resting state connectivity data.

| Contextualizing Variable | Parental Education $r_{cv}$ | Attenuation Magnitude | Attenuation $p$ value |
| --- | --- | --- | --- |
| No Contextualizing Variables | 0.247 | -- | -- |
| Enrichment Activities<sup>‡</sup> | 0.230 | 0.016 | 0.0439 |
| Family Support<sup>§</sup> | 0.247 | -0.001 | n.s. |
| School Support<sup>¶</sup> | 0.246 | 0.000 | n.s. |
| Child's Grades | 0.219 | 0.028 | 0.0001* |
| Child's General Cognitive Ability<sup>#</sup> | 0.189 | 0.058 | 0.0001* |
| All Contextualizing Variables | 0.175 | 0.071 | 0.0001* |

<sup>‡</sup> Intellectual/Cultural Orientation subscale of the Family Environment Scale
<sup>§</sup> Acceptance subscale of the Children's Report of Parental Behavior Inventory
<sup>¶</sup> School Environment subscale of the School Risk and Protective Factors Survey
<sup>#</sup> Latent Variable Created from ABCD Neurocognitive Battery
<sup>%</sup> Latent Variable Created from Child Behavior Checklist – Parent Report
\* Observed attenuation larger than all 10,000 in the permutation distribution.

## DISCUSSION

Using data from a large population-based sample of 5,821 9- and 10-year-olds in the ABCD Study, we evaluated associations between socioeconomic resources (SER) and youth functional connectomes using whole-brain, network-wise, and connection-wise approaches. Across these three levels of analysis, we observed widespread associations between SER and the developing connectome, with convergent evidence that parental education was the primary driver of these associations. Moreover, these parental education associations with the functional connectome were concentrated in somatosensory and subcortical networks, suggesting a spatial “footprint” for parental education effects that is somewhat distinct from other recently studied constructs (i.e., general cognitive ability and general psychopathology). Overall, these results add a new data-driven multivariate perspective on links between household SER and the child’s developing functional connectome. Moreover, they potentially illuminate a primary role for parental education in explaining how socioeconomic adversity gets “under the skin” to shape the developing brain.

Previous examinations of the associations between SER and functional connectivity patterns in the brain have largely relied on regionalist and apriorist methods (e.g., 31, 32), see (17) for a review. That is, SER-brain associations have been examined within individual pre-selected regions (e.g., amygdala – ventromedial prefrontal connectivity) based on prior theory. In addition, previous studies have generally been small, involving tens to hundreds of subjects (17), and these sample sizes may generally be too small for reliable statistical inference (19). Against the backdrop of this previous work, our study takes breaks new ground in taking a multivariate, brain-wide, data-driven approach. In whole-brain analysis, we found functional connectivity patterns across the entire connectome captured 9.0% of the variance in SER in held-out subjects, with statistically significant generalization of these SER-associated connectivity patterns to all 19 out of 19 held-out sites. In network-wise analysis, we found SER has distributed effects throughout the brain, with statistically significant effects observed at 77 of 120 cells (i.e., sets of connections linking large-scale networks). In connection-wise analysis, we found strong deviations between observed versus expected p-value distributions, with a notable pattern of “early lift-off” (see Figure 3), indicating the presence of highly distributed SER-related associations throughout the connectome.

Interestingly, our results are reminiscent of the distributed architecture of complex traits now recognized in genetics (37). There too, it was initially assumed that complex traits were represented by few genetic loci, each of large effect, leading to the popularity of candidate genes studies in which loci were selected for study based on prior theory. However, the polygenic nature of complex human traits is now considered the norm, wherein phenotypes result from the cumulative impact of hundreds of thousands of genetic variants, each of very small effect (38, 39). We here demonstrate a similar pattern in which SER-brain associations with the brain are analogously highly “poly-connectic”, implicating thousands of connections across the connectome, see also (40). Our results highlight the need for developmental neuroscience research on the effects of poverty on the brain to expand the toolkit of analysis methods beyond region-specific approaches to better capture what are likely to be highly distributed brain-wide effects (18, 41, 42).

We disentangled neural patterns of three indices of SER (parental education, household income-to-needs, and neighborhood disadvantage) using three different multivariate methods, each focused on different levels of analysis (whole-brain, network-wise, and connection-wise). These three methods convergently supported the conclusion that parental education has the most potent relationships with brain connectivity patterns in this large, population-based sample. Each of the three preceding method has its own strengths and weaknesses. For example, predictive modeling is optimized for aggregating widespread signals across the brain (40), but it is poor at localizing signals (43). Meanwhile, NCA is better for localizing effects but makes assumptions about network boundaries that may turn out to be suboptimal. Quantile-quantile modeling makes no assumptions about network boundaries but is not well suited for quantifying multivariate effect sizes. By using these three methods in combination, however, we address the limitations of each method and provide evidence of the robustness of our results across different analytic choices.

Parental education neural associations were preferentially concentrated in somatosensory/subcortical regions. This result agrees with previous seed-based studies that also found SER effects on connectivity patterns of amygdala, hippocampus, and striatum (35, 36, 44). The present study nonetheless goes beyond previous work in providing a brain-wide comparative perspective (i.e., parental education-associated connectivity is more concentrated in somatosensory/subcortical networks compared to other networks)—something that cannot be readily revealed by seed-based methods. Moreover, we showed this pattern of preferential somatosensory/subcortical concentration contrasts with general psychopathology (which is concentrated in default network and control networks; 11) and general cognitive ability (which is widespread across all networks; 12). Taken together, these results point to a distinctive spatial profile across the connectome of parental education that should be the focus of further investigation.

Several recent studies have also examined associations between SER and resting state functional connectivity in the same ABCD study dataset and reached somewhat different conclusions (45–47). These studies generally found much weaker brain-behavior associations than what is reported here. In addition, one study (46) found significant unique effects of neighborhood disadvantage, which were not observed here. Key differences in analytic approach may shed light on these divergent findings. The present study used connection-resolution connectomic data for each subject, encompassing 87,153 connections per connectome, as the input datatype for all analyses. In contrast, these other studies used summary statistics (available through the ABCD Data Exploration and Analysis Portal; https://deap.nimhda.org) in which each subject’s connectome is reduced to 78 numbers representing mean connectivity between each pair among 12 large-scale networks. In Supplement Figure S5 and Table S3, we report results from the current study alongside results that would have been seen were we to have used these summary statistics. We found that for SER, as well as for parental education, household income-to-needs, and neighborhood disadvantage, between 68 to 80% of the multivariate signal associated with these variables is lost when using the cell mean summary data (with 78 features per subject) rather than the connection resolution data (with 87,153 features per subject). It is likely that this signal loss, as well as other differences in analysis choices (discussed in Supplement §11), explain the weaker effect sizes observed in these studies and the different pattern of effects. Of note, the use of summary statistics in place of connection-level data has not been extensively validated, and the preceding results suggest the need for caution in adopting this approach.

One pathway through which SER (and particularly parental education) is thought to shape youth cognitive and socioemotional development is through the provision of cognitively-stimulating activities (48–50) that shape both cognitive skills (e.g., general cognitive abilities and academic skills (51–53) and non-cognitive skills including socioemotional skills (e.g., emotion regulation and impulse control). To better understand the proximal pathways through which parental education relates to whole-brain resting-state connectivity patterns, we conducted additional multivariate predictive modeling analyses with additional control variables. Home enrichment activities, child’s grades, and child’s cognitive abilities partially attenuated the multivariate association between parental education and resting-state connectivity patterns. This finding suggests there are rich, interwoven links between parental education, cognitive stimulation, and child cognitive/academic outcomes (34, 53). The full set of five contextualizing variables accounted for nearly 30% of the multivariate association between parental education and restingstate connectivity patterns. Regarding the remaining 70% of the effect, it is possible that the contextualizing variables that we assessed were measured imperfectly, and improved measurement would lead to larger attenuation. Alternatively, it is possible that other contextualizing variables that were not assessed at the current time in the ABCD dataset play an important role in the parental education-functional connectivity association.

One such variable is parental educational expectations, which are found to predict youth achievement longitudinally and over-and-above SER (54). Another may be genotype (55), wherein genetic variants shared by parent and child may account for some of the parental education-functional connectivity association (note: children’s genetic data are collected in the ABCD study but parents’ are not). A third possibility is that these connectivity patterns linked to parental education have behavioral sequalae at later ages (e.g., mid-to late adolescence), and thus they do not share variance with the contextualizing variables measured at ages 9 and 10. Overall, although the present study firmly establishes distributed functional connectivity patterns associated with levels of parental education, the specific pathways through which parental education becomes linked to brain connectivity await further elucidation (22).

This study has limitations and caution should be taken in interpreting its results. A key limitation is that this study uses cross-sectional data from the baseline wave of the ABCD study. This type of data cannot be used to infer causal relationships between modeled variables (56), and stronger inferences about “mediation” and/or causal relationships require other kinds of data, such as longitudinal data or experimental manipulations (56, 57). Additionally, the results of the current study should not be used to perpetuate *static*, *deficit* interpretations of development (58). The brain is a highly plastic organ with abundant capacities to learn/relearn, modify, and adjust, consistent with the observation that substantial neural change and reorganization extends through late adolescence up to young adulthood (59, 60). It is altogether possible that many of the SER-associated neural patterns we observed represent compensatory adjustments that help youth adaptively navigate features of their local environmental milieu (e.g., constrained opportunities, uncertainty) (61). It is also noteworthy that SER need not be static during childhood, and this includes parental education. A recent analysis (62) in a nationally-representative cohort reported that 17% of mothers completed additional education after the transition to motherhood. Moreover, this figure was further elevated (43%) among the most disadvantaged mothers, thus challenging assumptions about the “typical” sequence of life events. For parents who want to pursue additional schooling and/or certifications, policies and programs that provide childcare and workforce training can be effective (63). These observations point to future directions of research involving interventions on parental education levels and suggest potential downstream policy implications of the present results.

In sum, in a large, rigorously characterized sample of youth, we identified highly distributed, brain-wide functional connectivity patterns linked to SER. Moreover, we demonstrated that parental education was the primary driver of these associations, advancing our understanding of how socio-environmental factors are linked to the developing connectome.

## METHODS

### 1. Sample and Data

The ABCD study is a multisite longitudinal study with 11,875 children between 9-10 years of age from 22 sites across the United States. The study conforms to the rules and procedures of each site’s Institutional Review Board, and all participants provide informed consent (parents) or assent (children). Data for this study are from ABCD Release 3.0.

### 2. Data Acquisition, fMRI Preprocessing, and Connectome Generation

High spatial (2.4 mm isotropic) and temporal resolution (TR=800 ms) resting state fMRI was acquired in four separate runs (5min per run, 20 minutes total). Preprocessing was performed using fMRIPrep version 1.5.0 (64). Briefly, T1-weighted (T1w) and T2-weighted images were run through recon-all using FreeSurfer v6.0.1, spatially normalized, rigidly coregistered to the T1, motion corrected, normalized to standard space, and transformed to CIFTI space.

Connectomes were generated for each functional run using the Gordon 333 parcel atlas (65), augmented with parcels from high-resolution subcortical (66) and cerebellar (67) atlases. Volumes exceeding a framewise displacement (FD) threshold of 0.5mm were marked to be censored. Covariates were regressed out of the time series in a single step, including: linear trend, 24 motion parameters (original translations/rotations + derivatives + quadratics, aCompCorr 5 CSF and 5 WM components and ICA-AROMA aggressive components, high pass filtering at 0.008Hz, and censored volumes. Next, correlation matrices were calculated. Full details of preprocessing and connectome generation can be found in the Supplement as well as the automatically-generated FMRI Prep Supplement.

### 3. Inclusion/Exclusion

There are 11,875 subjects in the ABCD Release 3.0 dataset. Subjects were excluded for: failing ABCD QC, insufficient number of runs each 4 minutes or greater, failing visual QC of registrations and normalizations, and missing data required for regression modeling. This left us with N=5,821 subjects across 19 sites for the main sample analysis, and details of exclusions are provided in the Supplement.

### 4. Neuroimaging Analysis

To quantify brain-wide relationships between functional connectivity patterns and outcome variables of interest including SER, we used principal component regression (PCR) predictive modeling (26, 68) (see Figure S2). In brief, this method performs dimensionality reduction on resting state connectomes, fits a regression model on the resulting components, and applies this model out of sample in a leave-one-site-out cross-validation framework. To identify network-wise brain-behavior relationships, we used network contingency analysis (NCA) (27–29). In brief, for each cell (set of connections linking pairs of large-scale networks), this method identifies whether the count of connections significantly related to an outcome variable of interest exceeds what is expected by chance. To quantify connection-wise brain-behavior relationships, we used quantile-quantile modeling (30). In brief, we first calculated the p-value at each connection for the association between that connection and an outcome variable of interest. We then rank ordered these p-values and compared them to the rank-ordered distribution of p-values expected under the global null hypothesis.

In implementing the three preceding methods, we control for the effect of a number of nuisance covariates, specifically sex assigned at birth, self-reported race-ethnicity, age, age squared, mean FD and mean FD squared. For all three methods, we assessed statistical significance with nonparametric permutation tests, in which the procedure of Freedman and Lane (69) was used to account for covariates. In addition, exchangeability blocks were used to account for twin, family, and site structure and were entered into Permutation Analysis of Linear Models (PALM) (70) to produce permutation orderings. Details on all the preceding neuroimaging analyses are provided in the Supplement.

### 5. Latent Variable Modeling

We constructed a latent variable for socioeconomic resources by applying exploratory factor analysis to household income-to-needs, parental education, and neighborhood disadvantage. Household income-to-needs represents the ratio of a household’s income relative to its need based on family size, and details on its calculation are provided in the Supplement. Parental education was the average educational achievement of parents or caregivers. Neighborhood disadvantage scores reflect an ABCD consortium-supplied variable (reshist_addr1_adi_wsum). In brief, participant’s primary home address was used to generate Area Deprivation Index (ADI) values, which were weighted based on results from Kind *et al* (71) to create an aggregate measure.

The general psychopathology factor (P-factor) was produced from bifactor modeling of the parent-rated Child Behavior Checklist (CBCL) (72), and was described in detail in our previous studies in ABCD (73, 74). The general cognitive ability (GCA) variable was produced from bifactor modeling of the ABCD neurocognitive battery, and was also described in detail in our previous studies in ABCD (33, 73). Additional details on construction of the preceding latent variables are provided in the Supplement.

### 6. Code Availability

The ABCD data used in this report came from NDA Study 901, 10.15154/1520591, which can be found at https://nda.nih.gov/study.html?id=901. Code for running analyses can be found at https://github.com/SripadaLab/ABCD_Resting_Socioeconomic_Resources.

## Supporting information

Supplement FMRIPrep Methods

## ACKNOWLEDGEMENTS

Data used in the preparation of this article were obtained from the Adolescent Brain Cognitive Development (ABCD) Study (https://abcdstudy.org), held in the NIMH Data Archive (NDA). This is a multisite, longitudinal study designed to recruit more than 10,000 children age 9-10 and follow them over 10 years into early adulthood. The ABCD Study is supported by the National Institutes of Health and additional federal partners under award numbers U01DA041022, U01DA041028, U01DA041048, U01DA041089, U01DA041106, U01DA041117, U01DA041120, U01DA041134, U01DA041148, U01DA041156, U01DA041174, U24DA041123, and U24DA041147. A full list of supporters is available at https://abcdstudy.org/nih-collaborators. A listing of participating sites and a complete listing of the study investigators can be found at https://abcdstudy.org/principal-investigators.html. ABCD consortium investigators designed and implemented the study and/or provided data but did not necessarily participate in analysis or writing of this report. This manuscript reflects the views of the authors and may not reflect the opinions or views of the NIH or ABCD consortium investigators. The ABCD data repository grows and changes over time. The ABCD data used in this report came from NDA Study 721, 10.15154/1504041, which can be found at https://nda.nih.gov/study.html?id=721. NDA Study 901, 10.15154/1520591, which can be found at https://nda.nih.gov/study.html?id=901

This work was supported by the following grants from the United States National Institutes of Health, the National Institute on Drug Abuse, and the National Institute on Alcohol Abuse and Alcoholism: R01MH123458 (CS), U01DA041106 (CS, MH, LH), T32 AA007477 (KC), K23 DA051561 (AW). This research was supported in part through computational resources and services provided by Advanced Research Computing at the University of Michigan, Ann Arbor.

## Supplemental Methods and Results

### 2. Data Acquisition, fMRI Preprocessing, and Connectome Generation

Imaging protocols were harmonized across sites and scanners. High spatial (2.4 mm isotropic) and temporal resolution (TR=800 ms) resting state fMRI was acquired in four separate runs (5min per run, 20 minutes total). The entire data pipeline described below was run through automated scripts on the University of Michigan’s high-performance cluster, and is described below.

Preprocessing was performed using fMRIPrep version 1.5.0 (64), and detailed methods automatically generated by fRMIPrep software are provided in the fMRIPrep Supplement. T1-weighted (T1w) and T2-weighted images were run through recon-all using FreeSurfer v6.0.1. T1w images were also spatially normalized nonlinearly to MNI152NLin6Asym space using ANTs 2.2.0. Each functional run was corrected for fieldmap distortions, rigidly coregistered to the T1, motion corrected, and normalized to standard space. ICA-AROMA was run to generate aggressive noise regressors. Anatomical CompCor was run and the top 5 principal components of both CSF and white matter were retained. Functional data were transformed to CIFTI space using HCP’s Connectome Workbench. All preprocessed data were visually inspected at two separate stages to ensure only high-quality data was included: After co-registration of the functional data to the structural data and after registration of the functional data to MNI template space.

Connectomes were generated for each functional run using the Gordon 333 parcel atlas (65), augmented with parcels from high-resolution subcortical (66) and cerebellar (67) atlases. Volumes exceeding a framewise displacement threshold of 0.5mm were marked to be censored. Covariates were regressed out of the time series in a single step, including: linear trend, 24 motion parameters (original translations/rotations + derivatives + quadratics), aCompCorr 5 CSF and 5 WM components and ICA-AROMA aggressive components, high pass filtering at 0.008Hz, and censored volumes. Next, correlation matrices were calculated for each run. Each matrix was then Fisher r-to-z transformed, and then averaged across runs for each subject yielding their final connectome.

**Figure S1:**
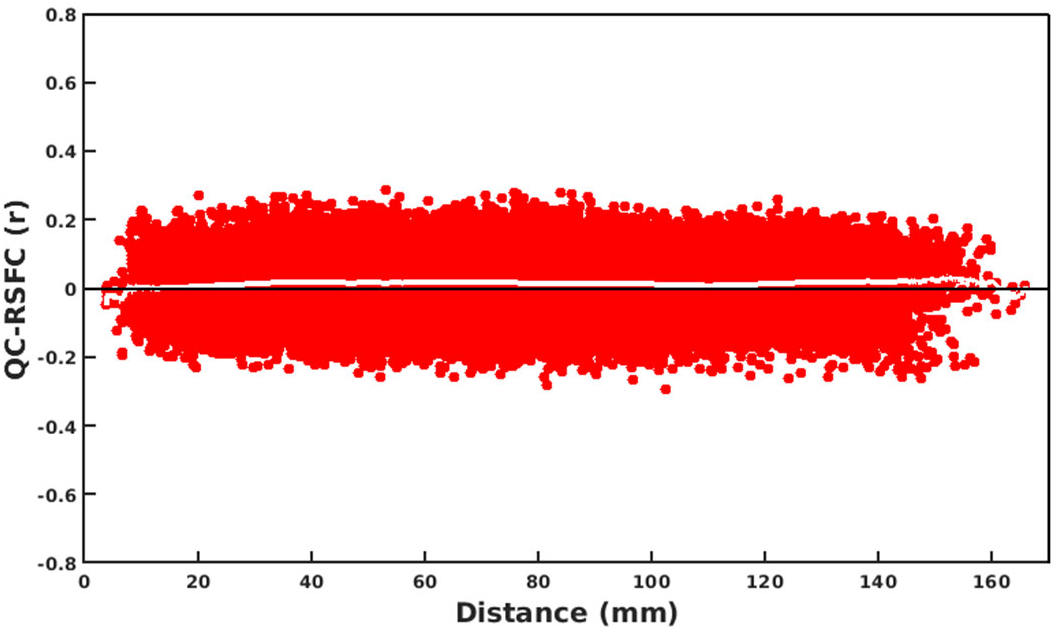
Quality Control-Resting State Functional Connectivity Plot.

We used multiple procedures listed above to limit the effect of head motion on resting state connectivity maps. To assess the effectiveness of these procedures, we produced a quality control resting state functional connectivity (QC-RSFC) plot (75, 76). This plot shows the relationship between mean framewise displacement and connectivity for edges binned by distance. Motion effects produce a sloped line (distance-dependent artifact), while a flat line is indicative of minimal motion-related effects. The RSFC-QC plot for our ABCD resting state data showed a flat line (Figure S1), providing additional evidence that our stringent motion correction strategies were effective.

### 3. Inclusion/Exclusion

There are 11,875 subjects in the ABCD Release 3.0 dataset. Screening was initially done using ABCD raw QC to limit to subjects with 2 or more good runs of resting data as well as a good T1 and T2 image (QC score, protocol compliance score, and complete all =1). This resulted in 9580 subjects with 2 or more runs that entered preprocessing. Each run was subsequently visually inspected for registration and warping quality, and only those subjects who still had 2 or more good runs were retained (N=8858). After connectome generation, runs were excluded if they had less than 4 minutes of uncensored data, and next subjects were retained only if they had 2 or more good runs (N=6568). Finally, subjects who were missing data required for SER factor modeling were dropped and sites with fewer than 75 subjects were dropped. This left us with N=5821 subjects across 19 sites for the neuroimaging analysis, and demographic characteristics of the overall and included sample are shown in Table S1.

**Table S1:**
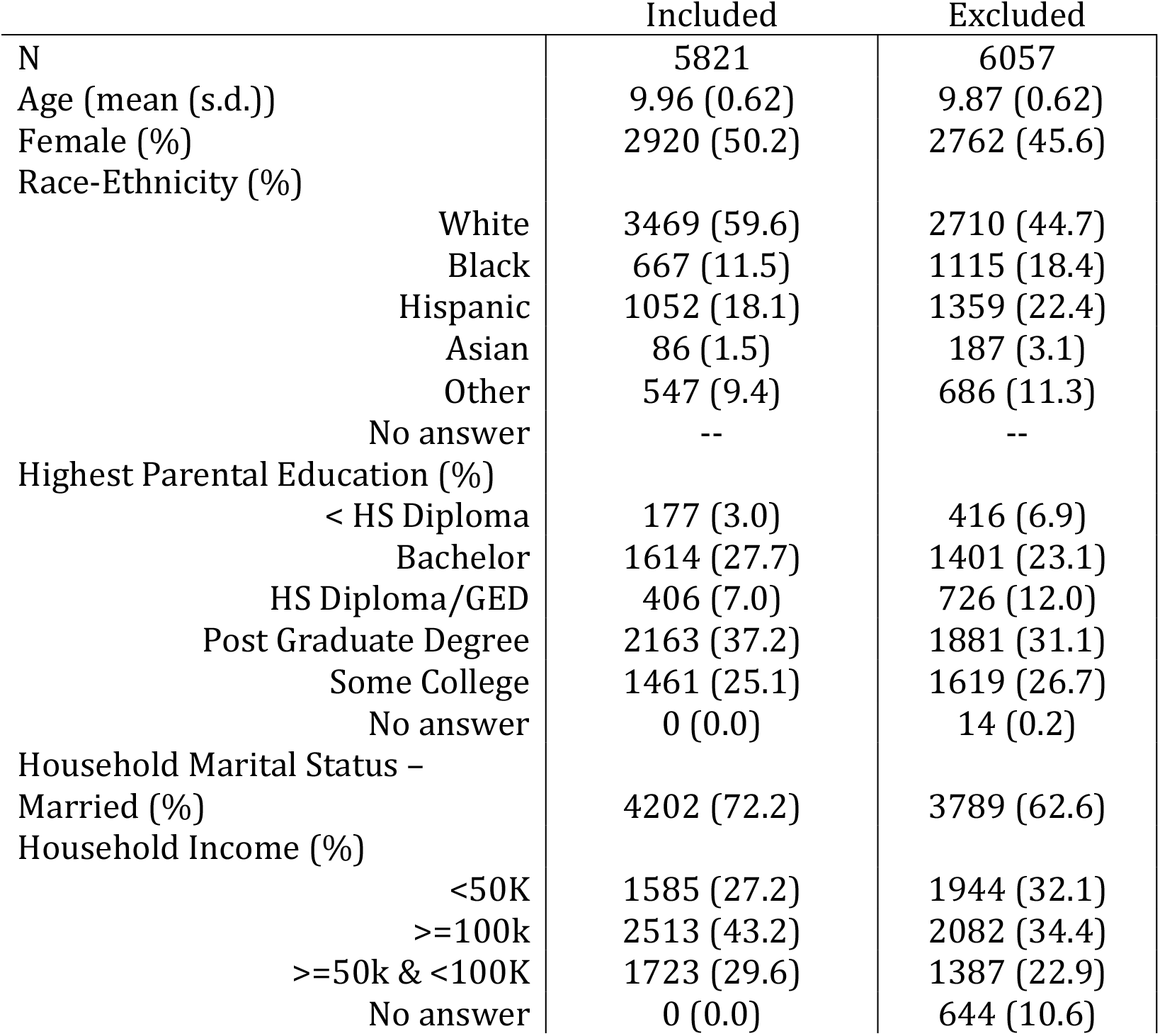
Demographic Characteristics of Included Versus Excluded Subjects.

|  | Included | Excluded |
| --- | --- | --- |
| N | 5821 | 6057 |
| Age (mean (s.d.)) | 9.96 (0.62) | 9.87 (0.62) |
| Female (%) | 2920 (50.2) | 2762 (45.6) |
| Race-Ethnicity (%) |  |  |
| White | 3469 (59.6) | 2710 (44.7) |
| Black | 667 (11.5) | 1115 (18.4) |
| Hispanic | 1052 (18.1) | 1359 (22.4) |
| Asian | 86 (1.5) | 187 (3.1) |
| Other | 547 (9.4) | 686 (11.3) |
| No answer | -- | -- |
| Highest Parental Education (%) |  |  |
| < HS Diploma | 177 (3.0) | 416 (6.9) |
| Bachelor | 1614 (27.7) | 1401 (23.1) |
| HS Diploma/GED | 406 (7.0) | 726 (12.0) |
| Post Graduate Degree | 2163 (37.2) | 1881 (31.1) |
| Some College | 1461 (25.1) | 1619 (26.7) |
| No answer | 0 (0.0) | 14 (0.2) |
| Household Marital Status – Married (%) | 4202 (72.2) | 3789 (62.6) |
| Household Income (%) |  |  |
| <50K | 1585 (27.2) | 1944 (32.1) |
| >=100k | 2513 (43.2) | 2082 (34.4) |
| >=50k & <100K | 1723 (29.6) | 1387 (22.9) |
| No answer | 0 (0.0) | 644 (10.6) |

### 4. Principal Components Regression-Based Multivariate Predictive Modeling

**Figure S2:**
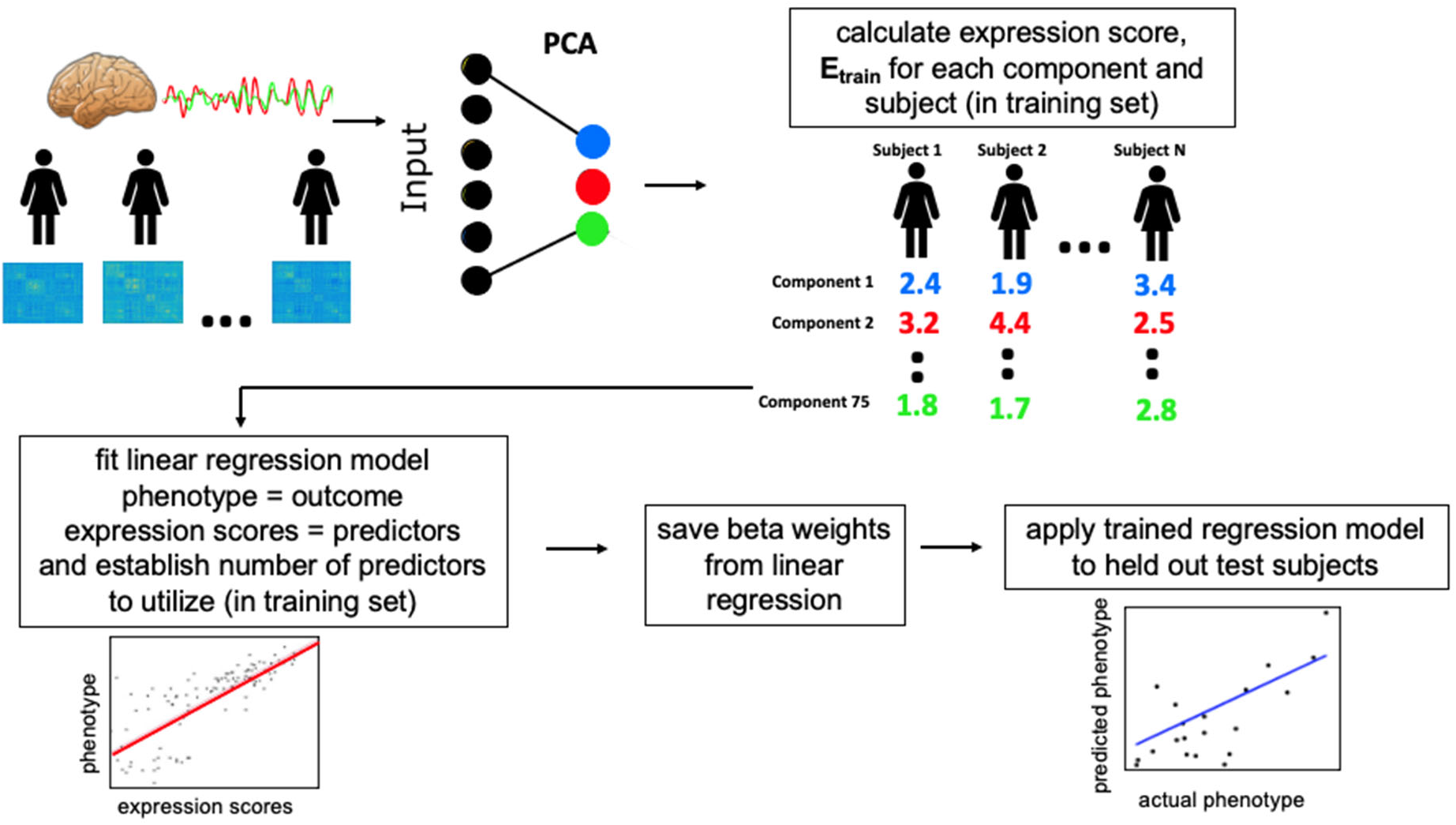
Steps of Principal Component Regression Predictive Modeling.

We implemented principal component regression (PCR) (77) as a multivariate predictive modeling method for identifying brain behavior relationships (26) (see Figure S2). The method involves two key steps: 1) Use principal components analysis (PCA) to find a set of components that capture *inter-individual* differences in brain features; 2) Use multiple regression in a crossvalidation framework to link expression scores for these components to phenotypes of interest. In previous work, we often used the more general name brain basis set (BBS) for this approach to capture commonalities with work by our group and others that use alternative methods for step 1 (e.g., independent component analysis (78, 79) or community detection (80, 81)). We chose the PCR approach for this study because our previous work showed it has high test-retest reliability (82) and predictive accuracy (68, 83), and generally performs as well or better than alternative methods such as support vector regression and ridge regression (82).

We performed PCA dimensionality reduction on an *n* subjects by *p* connectivity features matrix, yielding *n* principal components (i.e., directions in the feature space) that represent interindividual differences in connectivity. Per-subject expressions scores for a subset of *k* of these connectivity components then entered multiple regression modeling to identify linear associations with phenotypes of interest (here, GCA scores). Of note, we selected *k* using 5-fold cross validation within the training data, as in our previous work (82).

To assess accuracy and generalizability of PCR predictive models, we used leave-one-site-out cross-validation. In each fold of the cross-validation, data from one of the 19 sites served as the held-out test dataset and data from the other 18 sites served as the training dataset. Additionally, to ensure separation of train and test datasets, at each fold of the cross-validation, a new PCA was performed on connectomes in the training dataset, and expression scores of these brain components were calculated for the test set. Note that by employing leave-one-site-out, members of twinships and sibships are never present in both training and test samples. We assessed the performance of PCR predictive models with cross-validated Pearson’s correlation and cross-validated partial eta squared.

In each fold of the leave-one-site out cross-validation, PCR predictive models were trained in the train partition with the following covariates (unless explicitly stated otherwise for specific analyses): sex, race-ethnicity, age, age squared, mean FD and mean FD squared. To maintain strict separation between training and test datasets, regression coefficients for the covariates learned from the training sample were applied to the test sample to calculate effect size measures (Pearson’s correlation_cross-validated_ and R^2^_cross-validated_). This procedure is described in detail in our previous publications (33, 83).

We assessed the significance of all cross-validation-based correlations with non-parametric permutation tests. We randomly permuted the 5,821 subjects’ outcome variable values 10,000 times and reran the PCR predictive modeling stream at each iteration, yielding a null distribution of correlation values. The procedure of Freedman and Lane (69) was used to account for covariates. In addition, exchangeability blocks were used to account for twin, family, and site structure and were entered into Permutation Analysis of Linear Models (PALM) (70) to produce permutation orderings, as described in detail in the Supplement.

### 5. Network Contingency Analysis (NCA)

**Figure S3:**
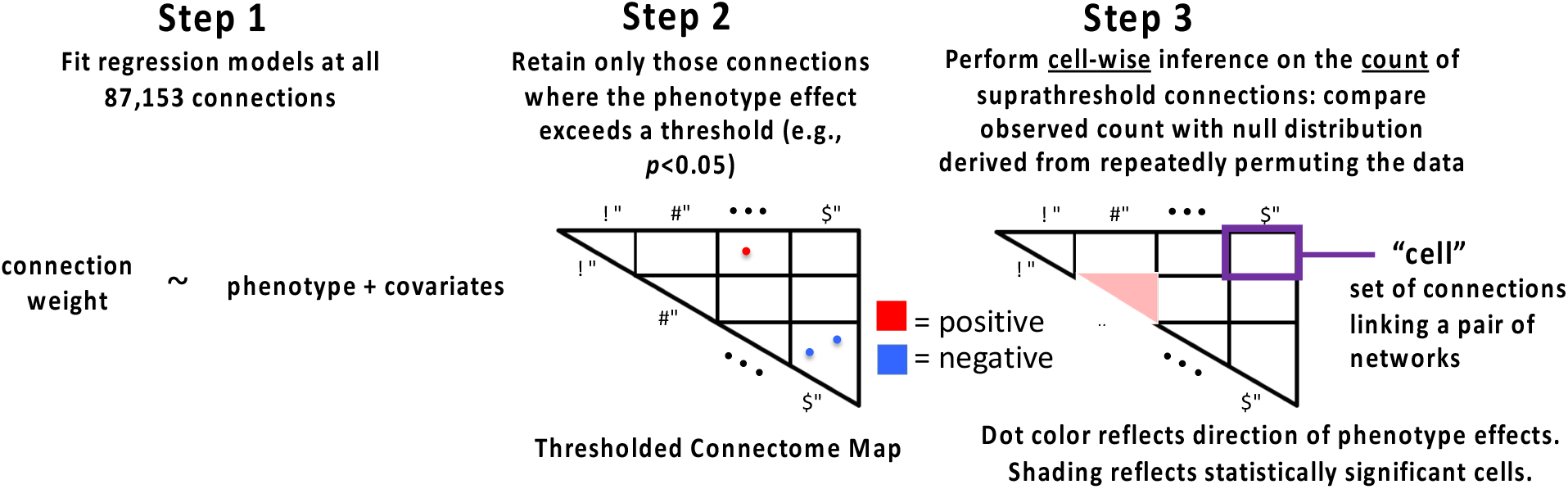
Steps for Network Contingency Analysis.

NCA is a count-based approach to quantifying altered connectivity (see Figure S3), see our prior work (27, 28, 84) for more details, and see also (29) for a related statistical treatment. In the current application, we applied the approach to “cells”; each cell is the set of connections linking a pair of the 16 networks in the Gordon parcellation augmented with additional networks (120 total cells). In Step 1 of the analysis, we fit a multiple regression model at each edge of the connectome with edge connectivity weight as the outcome variable and SER-related variables as the predictor of interest, while including sex, race-ethnicity, age, age^2^, mean FD, and mean FD^2^ as covariates. In step 2, we identified all connections in which the SER-related variable effect exceeds (is more significant than) a *p*<0.05 threshold (“NCA threshold”). In Step 3, we counted the suprathreshold connections separately for each cell, assessing whether this number exceeds the number that would be expected by chance alone. The distribution under chance was generated by non-parametric permutation tests (85). We randomly shuffled subjects’ edgewise connectivity weights 10,000 times (i.e., subject_i_’s edge weights were randomly switched with subject_j_’s) and recalculated the count of suprathreshold edges for each cell at each iteration. Permutation p-values were then calculated and corrected for multiple comparisons across 120 cells using the false discovery rate (86) with alpha set at p<0.05. The procedure of Freedman and Lane (69) was used to account for covariates. In addition, exchangeability blocks were used to account for twin, family, and site structure and were entered into Permutation Analysis of Linear Models (PALM) (70) to produce permutation orderings.

### 6. Quantile-Quantile Modeling

Quantile-quantile modeling is technique to simultaneously assess whether a collection of many statistical tests deviates from the expected null (30) and is commonly used in genome wide association studies (87). We adapt this approach to assess brain-wide associations by substituting genomic SNPs with edges from the connectome. In the first step, an association test is performed for each edge of the connectome by fitting a multiple regression model with the phenotype of interest as the outcome variable and edge connectivity weight as the predictor of interest, while including sex, race-ethnicity, age, age^2^, mean FD, and mean FD^2^ as covariates. In step 2, the observed p-values for the estimated effect size at each edge are then negative-log transformed, and then sorted from smallest to largest. In step 3, the rank-ordered negative-log transformed p-values are plotted versus negative-log transformed linearly scaled points on the x-axis - ie: the x-y coordinates for the plotted points are (−log(1), −log(largest p-value)), (−log(n-1/n), −log(2^nd^ largest p-value)),… (−log(1/n), −log(smallest p-value)). In step 4, the previous 3 steps are repeated in a permutation framework to generate the non-parametric null distribution. For the permutations, the procedure of Freedman and Lane [52] was used to account for covariates and exchangeability blocks were used to account for twin, family, and site structure and were entered into Permutation Analysis of Linear Models (PALM) [53] to produce permutation orderings. The mean from the permutation-based null distribution is plotted in red, and the 95^th^ percentile range of the permutation distribution is shaded in gray. If there is no association between the phenotype of interest and brain functional connections, the points corresponding to the observed edgewise association tests will fall within the 95^th^ percentile of the permutation null distribution.

### 7. Head Motion Robustness Tests

**Table S2:** Tests of Robustness of Parental Education Associations with Children’s Functional Connectomes. controlled = controlling for the effects of household income-to-needs and neighborhood disadvantage; r_cv_ = cross-validated out-of-sample correlation between actual scores and predicted scores using resting state connectivity data; * = observed correlation higher than all 10,000 correlations in the permutation distribution

| Subsample | parental education variable | $r_{cv}$ | $p_{PERM}$ | resampling test p-value |
| --- | --- | --- | --- | --- |
| low motion | uncontrolled | 0.24 | 0.0001* | 0.49 |
|  | controlled | 0.12 | 0.0001* | 0.29 |

To further assess potential confounding with head motion, we performed “resampling difference tests” (28) on a low motion subsample (with mean framewise displacement < 0.2 mm). First, we generated 10,000 random subsamples of the same size as the low motion subsample and performed multivariate predictive modeling for the parental education variable in each of these random subsamples. We next located the observed out-of-sample correlation for parental education in the actual low motion subsample in this resampling distribution. We declared the result significant if the observed correlation was below the 5th percentile, indicating that the observed correlation is lower than those from random subsamples of the same size than one should expect by chance.

### 8. Concentration Analysis

We performed a concentration analysis to assess whether cells significantly associated with outcome variables of interest (based on the previously described NCA analysis) were concentrated within three functional clusters. In the first step, all cells were assigned to three functional clusters: somatosensory/subcortical, default/control, other. In step 2, we calculated the concentration of significant cells within each cluster (our statistic of interest), defined as the average number of NCA significant cells for a functional cluster normalized by the total number of NCA significant cells across all three clusters. In step 3, the non-parametric permutation null distribution of this statistic of interest was constructed by repeatedly shuffling cell labels across the entire connectome and recomputing this statistic. The p-value of the observed concentration of NCA results per cluster was calculated as the proportion of the permutation distribution greater than observed.

### 9. Attenuation Analyses

We performed multivariate predictive modeling with a set of basic covariates (model 1) and a set of additional covariates (model 2), and we quantified the attenuation in out-of-sample predictive performance between the two models (attributable to the addition of these covariates). We assessed the statistical significance of this attenuation using bootstrap analysis. We resampled subjects with replacement within each ABCD site, then repeated the above multivariate predictive models, i.e., model 1 and model 2, with each bootstrapped sample, and calculated the magnitude of attenuation at each iteration. This was repeated 10,000 times yielding a distribution of bootstrapped attenuation values. The p-value of the observed attenuation value was calculated as the proportion of bootstrapped values that were less than 0 (i.e., indicating attenuation). Note for these bootstrapped analyses, we set the number of components for predictive modeling to be 400, i.e., approximately the mean selected for the observed data, to avoid computational demands from selecting component number by nested cross-validation in each of 10,000 bootstrap iterations.

### 10. Latent Variable Modeling for Socioeconomic Resources, General Psychopathology, and General Cognitive Ability

**Figure S4:**
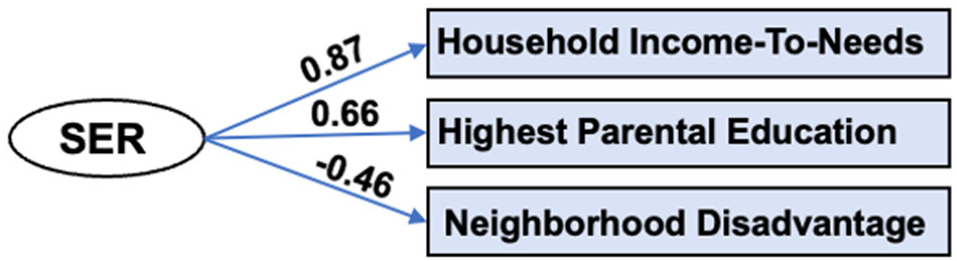
Factor Model of Socioeconomic Resources. Path estimates reflect standardized factor loadings.

We constructed a latent variable for socioeconomic resources by applying factor analysis to household income-to-needs, highest parental education, and neighborhood disadvantage in 10,578 participants that had all three variables (see Figure S4). The latent variable accounted for 47% of the variance in the observed scores. Household Income-to-Needs combines information on household income, poverty lines, and family size. Household Income covered all sources of income for family members, including wages, benefits, child support payments, and others. It was assessed in bins as follows: 1 <5,000, 2 5,000 - 11,999, 3 12,000 - 15,999, 4 16,000 - 24,999, 5 25,000 - 34,999, 6 35,000 - 49,999, 7 50,000 - 74,999, 8 75,000 - 99,999, 9 100,000 - 199,999, 10 More than 200,000, and we assigned each subject the midpoint for their bin. Household size was calculated from the ABCD Parent Demographic Survey. The poverty line was calculated for a household of that size based on the 2021 US Poverty Guidelines ($8,340 + $4,540 per person in the household). Finally, Household Income-to-Needs was calculated as the ratio of the combined income midpoint over the poverty line. Highest Parental Educational was the highest educational achievement by either parent or caregiver. Neighborhood disadvantage scores reflect an ABCD consortium-supplied variable (reshist_addr1_adi_wsum). Participant’s primary home address was used to generate Area Deprivation Index (ADI) values, which were weighted based on results from Kind *et al* (71) to create an aggregate measure. Higher scores on the factor indicate greater neighborhood disadvantage including higher percent of families living in poverty, increased unemployment, and lower levels of educational attainment at the neighborhood level.

#### General Psychopathology Factor

The general psychopathology factor (P-factor) used here is based on the parent-rated Child Behavior Checklist (CBCL) (72), age 6 to 18 form. A bifactor model was fit to eight CBCL scales, with a general P factor that all scales loaded onto (average scale loading = .69) and two specific factors. Further details as well as the factor model are provided in our previous publication in ABCD (73, 74).

#### General Cognitive Ability Factor

The general cognitive ability (GCA) factor used here is based on latent variable modeling of the ABCD Neurocognitive Battery. We used exploratory factor analysis and parallel analysis to arrive a three-factor solution. A subsequent bifactor model showed very good fit by conventional standards (χ^2^ (34)=443.16, *p*<0.001, RMSEA=0.03, CFI=0.99, TLI=0.98, SRMR=0.02), with the general factor capturing 75% of the variation in task scores [coefficient ω hierarchical (88)], and three domain-specific factors together accounting for 13% of the variation in task scores. Further details as well as the factor model are provided in our previous publication in ABCD (33).

### 11. Comparison of Connection Resolution Results with Results Using Cell Mean Summary Statistics

**Figure S5:**
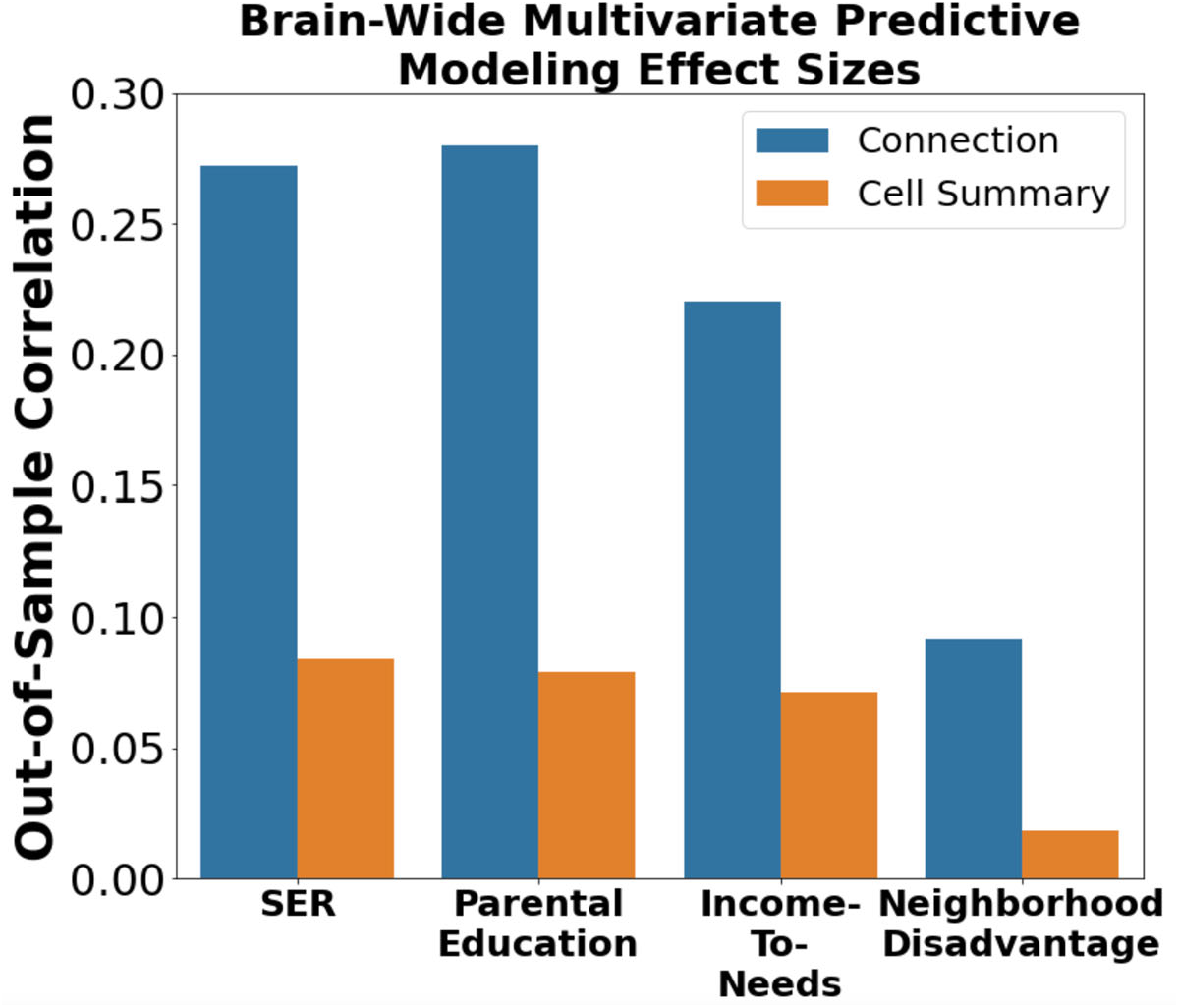
Effect Sizes for Connection Resolution Results Compared to Cell Mean Resolution Results. We compared multivariate predictive modeling analysis with connection resolution data (with 87,153 features per subject) and cell mean resolution data (with 78 features per subject). We found between 68 to 80% of the multi-variate signal associated with SER-related variables is lost when using the cell mean summary data rather than connection-resolution data.

**Table S3:** Effect Sizes and Percent Reduction in Effect Size for Connection Resolution Results Compared to Cell Mean Results. r_cv_ = cross-validated out-of-sample correlation between actual scores and predicted scores using resting state connectivity data.

| Variable† | Out-of-Sample $r_{cv}$ | | |
| --- | --- | --- | --- |
|  | CONNECTION | CELL SUMMARY | % Reduction |
| SER | 0.272 | 0.079 | 71 |
| Parental Education | 0.280 | 0.084 | 70 |
| Household Income-To-Needs | 0.220 | 0.071 | 68 |
| Neighborhood Disadvantage | 0.092 | 0.018 | 80 |

All analyses reported in the Main Manuscript were performed on whole-brain connectomes, each with 87,153 connections. Recent studies (45–47) of SER and SER-related variables in ABCD use summary statistics in which each subject’s connectome is reduced to 78 numbers representing mean connectivity between each pair among 12 large-scale networks, and these summary statistics are available through the ABCD Data Exploration and Analysis Portal; https://deap.nimhda.org. We repeated our multivariate predictive modeling approach with summary statistics data to assess whether the underlying signal associated with SER-related variables is preserved in the summary data. That is, we performed the same principal component regression with leave-one-site-out cross-validation described in §4 of Supplemental Methods, this time with 78 summary features per subject rather than 87,153 connections per subject. As shown in Figure S5 and Table S2, we found that between 68 to 80% of the multivariate signal associated with SER-related variables is lost when using the cell mean summary data rather than connection-resolution data. It is possible that this sizable reduction in signal explains the weaker effect sizes observed in these recent studies (45–47) as well as the different pattern of effects observed. Of note, there were in addition other differences in the current study that could contribute to differing results. We performed ICA AROMA to further reduce motion artifacts (89). In addition, we used a minimum post-censoring scan length threshold, 8 minutes based on (19), due to less reliable connectivity estimates with shorter scan lengths (90).

